# Dynamic gene gain and loss during the divergence of obligate biotrophic powdery mildew pathogens

**DOI:** 10.64898/2026.06.05.730400

**Authors:** Thomas C. Heaven, Helen M. Cockerton, Xiangming Xu, Matthew R. Goddard, Andrew D. Armitage

## Abstract

Powdery mildew outbreaks can result in devastating crop losses across both horticultural and cereal crops. The obligate biotrophic fungi responsible (Erysiphaceae) depend entirely on living plant hosts for survival, extracting nutrients exclusively from living host tissue. Despite their impact, genomic resources for dicot-infecting powdery mildew fungi remain limited, restricting understanding of differentiation in the effector complements that underpin biotrophy across mildew lineages. To address this, we sequenced and assembled genomes from three independent samples each of *Podosphaera leucotricha* (apple mildew) and *P. aphanis* sensu lato (strawberry & raspberry mildew). These novel genomes were analysed alongside 44 publicly-available Erysiphaceae genomes to reveal striking inter-species differences in effectorome complements. *P. leucotricha* encoded the largest predicted effector complement of any powdery mildew pathogen sequenced to date, with major expansions in RALPH and EKA effector families. By contrast, the closely related *P. aphanis* carried a comparatively small effector repertoire. Together, these data challenge the prevailing view that effector expansion is restricted to monocot-infecting mildews. We observed that the divergence of powdery mildew genera is associated with the contraction of gene families, consistent with a stepwise loss of genes. This pattern was also reflected in the loss of genes that are typically conserved across ascomycete fungi. Taken together our findings highlight the contrasting impact that biotrophic adaptation has had on pathogen genomes, namely the convergent loss of conserved fungal genes alongside the diversification of pathogen effectoromes in response to host immune landscapes.

## INTRODUCTION

Obligate biotrophic species are locked in a co-evolutionary arms race with their hosts. Their successful infection of plants depends on their repertoire of secreted effector proteins, which enable suppression of the host plant’s immune system (1–3). Members of the Erysiphaceae family, known as powdery mildew, are the obligate biotrophs responsible for powdery mildew diseases. These pathogens threaten the production of many economically important crops (4,5). For example, *Podosphaera* species infecting apple, strawberry, and cucurbits can cause substantial yield and quality losses, sometimes exceeding 30% (6–8).

The roles and evolutionary pressures acting on effectors differ markedly between fungi with different life histories. In necrotrophs and hemibiotrophs, effectors often function as toxins or cell-death inducers, with gene families expanding alongside secondary metabolite clusters and plant cell wall–degrading enzymes. By contrast, in obligate biotrophs, whose lifestyles depend entirely on keeping host cells alive while extracting nutrients, effectors must facilitate ongoing manipulation of living host tissue without triggering further defence responses. Biotroph effectors are therefore pivotal in determining host range. Consistent with their lifestyle, obligate biotrophs exhibit extensive genomic adaptations, including the convergent loss of primary metabolic genes and specialised repertoires of secreted effectors. Powdery mildew pathogens provide a compelling system for examining how such pressures shape effector evolution. The Erysiphaceae family forms a distinct monophyletic group, thus the most parsimonious explanation for their biotrophic lifestyle is descent from a common pathogenic ancestor (5,9–12). However, through divergence and adaptation, the family has since radiated into a diverse array of host-specific species. Powdery mildew effector proteins have been shown to target a range of host defence-related proteins and processes, such as hydrogen peroxide production, papillae formation, haustorial encasement, and programmed cell death (13–20). The ability of different Erysiphaceae species to infect different hosts has been associated with the differential arsenal of effectors they carry (21).

Effector paralogues have been shown to rapidly lose sequence similarity through diversification (22,23). The DNA sequences underling effectors are therefore highly diverse, and their identification relies upon very broad criteria (24,24). Effectors are typically small, secreted, lack known functional domains, and rarely have homologues in unrelated species (25,26). This is true of powdery mildew effectors (18,27). Reflecting this, fewer than 5.1% of *Blumeria* effector families have non-mildew leotiomycete homologues (28), and only a single effector orthogroup is shared across *Blumeria* spp.*, E. quercicola* and *E. necator* (29). In contrast, most non-effector Erysiphaceae genes have homologues in other ascomycetes (92% of total genes in *E. necator*) (30). As such, identifying secreted proteins that lack conservation outside of the Erysiphaceae is an important trait when defining and identifying Candidate Secreted Effector Proteins (CSEPs) of powdery mildew pathogens.

Monocot infecting *Blumeria* spp. are the most extensively studied powdery mildew pathogens to date (1,31). However, investigations of other genera have indicated that dicot infecting powdery mildew genera carry far fewer candidate secreted effector proteins than their monocot counterparts (29,32). Wu et al. (2018) identified four times as many genes encoding CSEPs in monocot (*Blumeria graminis* and *Blumeria hordei*) versus dicot powdery mildew species (*E. necator*, *Erysiphe neolycopersici* and *Golovinomyces cichoracearum*). Whether CSEP expansion is a unique feature of the *Blumeria* lineage, reflects strict host-specificity, or is driven by the prevalence of host resistance genes (*R* genes) remains unresolved. The application of comparative genomics across a broader range of Erysiphaceae species is required to elucidate the evolution of effector diversification and host range (29).

Within the *Podosphaera* genus, *Podosphaera leucotricha* only affects trees of the Roseascea family, primarily affecting apple, but also infecting almond, peach, pear, quince, and medlar (33,34). The *P. aphanis* species complex demonstrates still greater specificity, being limited to strawberry and *Rubus* crops (shrubs also in the Roseascea family) (35). The lack of genomic resources for the *Podosphaera* genus has hindered our ability to ask fundamental questions about the evolution of obligate biotrophy in horticultural powdery mildew pathogens. By sequencing and comparing multiple assemblies of *P. aphanis* s. lat. (strawberry & raspberry mildew) and *P. leucotricha* (apple mildew), this study tests the hypothesis that effectorome expansion is unique to monocot-infecting powdery mildew pathogens and examines how effector diversity contributes to host specificity. We further explore how effector gain, together with convergent loss of conserved fungal genes, shapes adaptation to obligate biotrophy. These comparative analyses advance our understanding of host specialisation in phytopathogenic fungi and reveal broader evolutionary principles governing biotrophic lifestyles.

## MATERIAL AND METHODS

### Mildew sampling

Samples were collected from naturally occurring powdery mildew outbreaks at Niab, East Malling, Kent, UK, 51.2859° N, 0.4533° E, across three growing seasons. *P. aphanis* samples were collected: from infected leaf material of *Fragaria* × *ananassa* ‘Malling Centenary’ plants growing under polytunnel conditions in May 2020, from infected fruit and stolons of *F. ananassa* ‘Malling Ace’ plants growing in polytunnel conditions in July 2021, and from leaves of a *Rubus* × *idaeus* population growing in field conditions in July 2020. *P. leucotricha* samples were collected: from the leaves of established *M. domestica* trees growing in exposed orchard conditions in April 2019 (varieties: ‘M1 I’; ‘Idared’; ‘Loopspy’; ‘M18’; ‘MM106’; ‘Gala’; ‘Jonathan’ - combined), June 2021 (varieties: ‘Northern Spy’; ‘M1’; ‘M9’; ‘Loopspy’; ‘Normandee’; ‘Granny Smith’; ‘Ottawa 3’; ‘Saturn’; ‘Pineapple Russet’; ‘M54-1’ - combined), and from leaves of a population of one year old *Malus* × *domestica* seedlings growing in glasshouse conditions in April 2020.

### DNA extraction, assembly and RNA-guided gene prediction

DNA/RNA extraction and genome assembly and gene prediction were performed for all six samples using the methodology described in Heaven et al. (2023) (S1) (36). Sequencing was performed on an Illumina NovaSeq platform using PE150 chemistry. RNA-seq reads from samples DRCT72020 and P112020 were utilised as evidence for gene models for *P. aphanis* and *P. leucotricha* genomes, respectively. Gene prediction was first performed by BRAKER v1.9 (13,989 - 16,533 genes) before supplementing with CodingQuarry predictions in intergenic regions between BRAKER gene models (1,077 - 3,203 genes per genome). Intergenic distances were extracted for comparison via a custom python script (S1).

### Reannotation of 44 published mildew genomes

Publicly available powdery mildew genomes and in-house assemblies were subject to standardised gene-prediction for consistent comparison across 50 total genomes. These gene predictions were performed as follows: genome assemblies were processed via the NCBI Foreign Contamination Screen (FCS) tool and assessed via Kraken 2 prior to gene annotation of ‘clean’ genome sequences (37–39); intron hint files were generated via the ProtHint v2.6.0 pipeline, using data from the fungal OrthoDB database (https://v100.orthodb.org/download/odb10_fungi_fasta.tar.gz, accessed on 14 August 2022) (40). BRAKER v1.9 was then run with the –epmode flag to perform gene prediction with hints as well as the – ab_initio flag to perform *ab initio* gene predictions without hints (41); gene duplication within genomes was assessed via BUSCO v4.0.6 analysis (42).

### Candidate Effector Identification

#### Candidate Secreted Effector Protein Prediction

Proteins were considered to be canonically secreted if they were predicted to contain <2 transmembrane domains and a signal peptide by any of the tools in the Predector v1.2.6 pipeline (43). CSEPs were defined as proteins predicted to be canonically secreted and lacking BLASTP (E ≤ 1 × 10⁻⁵) homology to *Podospora anserina* (GCF_000226545.1) or *Neurospora crassa* (GCF_000182925.2). Proteins additionally identified as effectors by EffectorP v3.0 were designated EffectorP CSEPs (44).

#### Effector Homologues

The Predector pipeline was used to search a curated set of experimentally validated fungal effector Hidden Markov Models (HMMs) (https://doi.org/10.6084/m9.figshare.16973665) including RALPH (RNAse-Like Proteins associated with the Haustoria) and EKA (effectors homologous to *AVR*_k1_ and *AVR*_a10_) genes from powdery mildew pathogens [downloaded 30/07/2022, https://doi.org/10.6084/m9.figshare.16973665], PHI-base v4.13, and the Pfam database [downloaded 30/07/2022] (43,45–48).

### Comparative analysis of conserved and unique pathogenicity factors in mildew species

#### Cross-Clade Erysiphaceae Orthology Analysis

Orthology among protein predictions from erysiphoid clade assemblies (Erysiphaceae and *Arachnopeziza araneosa*) was inferred using OrthoFinder v2.5.4 (S2, S3) (49,50). Three proteomes (*Erysiphe alphitoides* CLCBIO_7.0.3; *E. necator* GCA_016906895.1; *Golovinomyces magnicellulatus* GCA_006912115.1) were excluded due to suspected contamination. Only the longest isoform per gene was retained.

#### *Podosphaera aphanis* and *Podosphaera leucotricha* High-Confidence Gene Prediction Orthology

Orthologues among predicted proteins from sequenced genomes were identified using OrthoFinder v2.5.4 (49). Core orthogroups shared across all six assemblies, and species-exclusive orthogroups present in all *P. leucotricha* but absent from *P. aphanis* assemblies, or vice versa, were identified. The analysis was also repeated excluding the raspberry powdery mildew *P. aphanis* assembly (SCOTT2020). Venn diagrams were generated in R using ggvenn v0.1.9 (51). Exact 95% binomial confidence intervals for proportions of species-exclusive orthogroups were calculated using the Clopper-Pearson method.

EffectorP3-predicted CSEPs belonging to orthogroups exclusive to *P. aphanis* or *P. leucotricha* were examined as species-specific effector candidates. Each candidate protein was queried using TBLASTN v2.12.0 (52) against 44 publicly available mildew genomes (S2) and the six genomes generated in this study. The “-max_intron_length” parameter was set to the greater of 40 bp or 110% of the maximum intron length predicted for *P. aphanis* or *P. leucotricha* gene models. Hits with ≥ 70% identity across ≥ 70% of the largest predicted exon and E < 1 × 10^−20^ were retained. To identify cases where hits corresponded to distinct exons of the same gene, alignments were inspected in Geneious v2023.1.3 using Clustal Omega v1.2.2. Additional BLASTP v2.12.0 (52) searches were conducted against the six *P. aphanis* and *P. leucotricha* proteomes using the same thresholds.

### Host-pathogen co-evolutionary analysis

A phylogeny was constructed from the six newly generated assemblies and 44 publicly available Erysiphaceae genomes (S2). A total of 3,232 conserved single copy orthologous BUSCO genes were used for alignment and tree inference with IQ-TREE v2.3.0 (S1). Gene family expansions and contractions were then assessed using CAFE v5.1 based on orthogroups identified by the cross-clade Erysiphaceae orthology analysis (53). Following Mendes et al. (2021), gene families containing >100 members in any species were excluded. To account for assembly errors, an error model was estimated before final birth–death (λ) parameter estimation.

Gene loss of 99 Missing Conserved Ascomycete Genes (MCAGs) previously reported across mildews (31) was reassessed using BLAST-based searches across six genomes generated in this study, 44 publicly available Erysiphaceae genomes (S2) and 28 non-mildew controls (S3). A BLASTP database was built from predicted protein sequences of mildew and non-mildew fungi. Each of the 99 MCAGs defined by Spanu et al. (2010) was queried using BLASTP v2.12.0 with cuttoff E < 1 × 10⁻⁵ (52). Results were visualized in iTOL v5 (54). To validate positive hits, proteins showing MCAG identity were reciprocally searched against the *Saccharomyces cerevisiae* proteome and against UniProt (downloaded 1 October 2023) using DIAMOND v0.9.29 (55). Proteins returning hits to distantly related taxa were considered likely contaminants and excluded from further analysis.

## RESULTS

### Isolation of high-confidence *Podosphaera* assemblies through environmental sampling followed by bioinformatic filtration

To investigate the genomics of host adaptation within the *Podosphaera* clade and to place these findings in the broader context of evolution within the Erysiphaceae, six high-coverage genome assemblies (∼72.5 - 432×) were generated for *P. leucotricha* and *P. aphanis*, representing replicated samples within the *Podosphaera clade* (Table 1; Heaven et al., 2023). These assemblies were generated from conidial samples collected from natural infections, with powdery mildew pathogen sequences successfully recovered from mixed epiphytic datasets through contig-filtering, as applied in Heaven et al. (2023) (S1).

**Table 1:** Genome statistics for the *Podosphaera* genomes generated in this study: N50 represents the contig size of which 50% of the bases are in contigs of this size or greater. BUSCO- Benchmarking Universal Single-Copy Orthologues.

|  | *P. leucotricha* |  |  | *P. aphanis* |  |  |
| --- | --- | --- | --- | --- | --- | --- |
| ID | OGB2019 | P112020 | OGB2021 | DRCT72020 | DRCT72021 | SCOTT2020 |
| Host common name | apple | apple | apple | strawberry | strawberry | raspberry |
| Raw reads | 206,353,020 | 121,174,496 | 227,697,160 | 698,565,358 | 251,729,842 | 211,142,728 |
| Trimmed reads | 203,585,109 | 117,214,493 | 224,546,425 | 685,445,771 | 249,134,222 | 208,078,353 |
| Assembly size | 52,561,395 | 49,079,051 | 49,818,523 | 55,613,046 | 54,152,750 | 51,418,734 |
| Contigs >500bp | 10,456 | 6,805 | 7,325 | 12,359 | 8,198 | 10,355 |
| Largest conig | 136,346 | 100,493 | 108,534 | 77,136 | 559,432 | 75,917 |
| GC% | 41.71 | 41.76 | 41.54 | 43.06 | 43.62 | 43.06 |
| N50 | 19,114 | 20,168 | 19,522 | 11,409 | 12,165 | 8,889 |
| Median depth | 72.5 | 123.3 | 77.5 | 432 | 74.4 | 84.5 |
| BUSCO complete | 2,985 (92.3%) | 2,982 (92.2%) | 2,981 (92.2%) | 2,973 (91.9%) | 2,987 (92.4%) | 2,988 (92.2%) |
| Busco duplicated | 29 (0.9%) | 13 (0.4%) | 12 (0.4%) | 11 (0.4%) | 89 (2.8%) | 13 (0.4%) |
| BUSCO missing | 162 (5.0%) | 166 (5.1%) | 166 (5.1%) | 177 (5.5%) | 167 (5.1%) | 168(5.2%) |

Assembly sizes were 50.7 and 55.6 Mb for strawberry-infecting *P. aphanis*, 51.1 Mb for raspberry-infecting *P. aphanis*, and 48.9 – 49.4 Mb for *P. leucotricha*. BUSCO analysis indicated that all assemblies were of high quality, with 91.9 - 92.4% of 3,234 Leotiomycete orthologues identified as complete and single copy (Table 1). This level of completeness, comparable to that reported for other mildew genomes (90.9 - 94.8%), demonstrates that the gene space is well represented across all *Podosphaera* assemblies. Genome size was estimated at ∼160 Mb for both species based on the distribution of k-mer counts. The discrepancy between estimated genome size and assembly span reflects the exceptionally high repeat content typical of powdery mildew species (31,56,57). RepeatMasker identified 23 – 29 Mb of masked sequence per assembly, suggesting that up to 85% of the *Podosphaera* genomes are comprised of repetitive elements. In line with this, assembly contiguity (N50: 8,869–20,226 bp; contigs: 6,752–12,357; Table 1) compared favourably with previous short-read assemblies but was lower than that achieved by recent hybrid assemblies incorporating long-read data (28,31,58–61).

We analysed gene-to-gene and gene-to-transposable element (TE) distances in *P. leucotricha* and *P. aphanis*. These analyses revealed no extended gene-rich or gene-sparse regions, and only minimal differences in the spatial arrangement of genes with predicted housekeeping and effector functions (S4, S5). Average three prime intergenic distance was significantly greater than average five prime intergenic distance for BUSCO genes across all six assemblies (p = 8.95 × 10-6 to 8.79 × 10^−16^) (S6). In contrast, the average five prime intergenic distance was significantly greater than average three prime intergenic distance for RALPH and EKA genes in all six assemblies (p = 0.044 to 9.14 × 10^−4^). This may reflect more complex regulation of putative effector genes, as five prime regions typically contain regulatory elements controlling gene expression. However, differences in total intergenic distance between CSEP and BUSCO were small; mean total intergenic distances (3’ + 5’) for CSEP genes ranged from 3,226 to 3,913 bp, versus 2,746 to 3,522 bp for BUSCO genes (S6). Together, these results describe highly repetitive ‘one-speed’ genomes characteristic of powdery mildew fungi in which both conserved and lineage-specific genes are interspersed throughout a more-or-less uniformly repeat-rich landscape. The overall architecture of the *Podosphaera* genomes thus reinforces the view that genome expansion through TE proliferation, rather than compartmentalization, underlies much of the structural evolution observed across powdery mildew fungi.

### Multi-locus phylogenetics highlights differences in host range and evolutionary strategy between mildew genera

To investigate pathogen-host coevolution within the Erysiphaceae we reconstructed a concatenated phylogeny from 3,232 conserved single copy orthologous genes across the six newly assembled *Podosphaera* genomes, together with 44 publicly available genomes from the wider mildew clade (Figure 1). *P. leucotricha* formed a well-supported monophyletic group distinct from the *P. aphanis* complex, confirming deep divergence between apple-infecting and strawberry/raspberry-infecting species. The *P. aphanis* complex was recovered as a sister taxon to *P. cerasi* infecting cherry, consistent with shared ancestry among Rosales-associated powdery mildew species. Within the *P. aphanis* complex, analysis of internal transcribed spacer (ITS) and 28S regions extracted from our *de novo P. aphanis* s. lat. assemblies placed the raspberry sample within the recently proposed *P. ruborum* clade of *P. aphanis* and the two strawberry samples within the recently proposed *P. fragariae* clade, consistent with host-associated divergence.

**Fig. 1.**
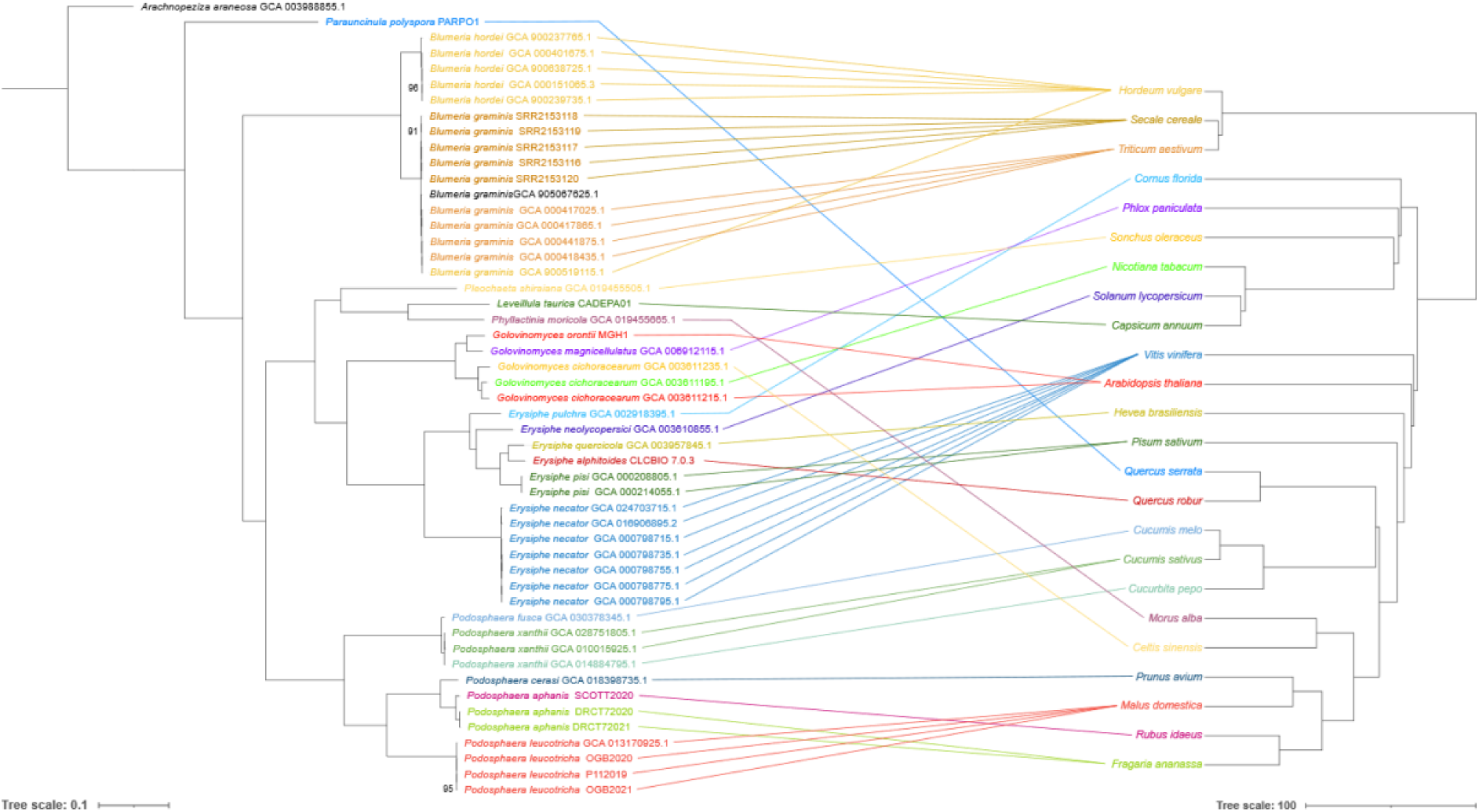
Phylogeny of sequenced powdery mildew pathogens and their hosts: Maximum likelihood phylogenetic trees are presented for Erysiphaceae genomes (3,232 orthologues) and host plants (425 orthologues). Powdery mildew pathogen species are colour-coordinated with, and connected to, the respective host plant from which they were sampled. Support values are shown for nodes with support <1.

Parallel reconstruction of Erysiphaceae host phylogenies from available plant genomes (Figure 1) revealed that phylogenetic relationships among *Podosphaera* pathogens were largely congruent with those of their respective host plants, consistent with a history of long-term host tracking and coevolution within this genus. In contrast, relationships in other mildew genera were not mirrored by those of their host taxa. For example, *Golovinomyces* spp., which have broad host ranges, can infect diverse host plants, whilst the endophytic group pathogens *Phyllactinia moricola* and *Leveillula taurica* grouped together but infect *Morus alba* (order Rosales) and *Capsicum annuum* (order Solanales), respectively.

Together, these results indicate that evolutionary trajectories within the Erysiphaceae clade are shaped by both conserved biotrophic specialisation and occasional host-jump events that promote diversification and ecological breadth across the Erysiphaceae. These findings reflect evolutionary dynamics across obligate biotrophs, wherein stable coevolutionary relationships coexist with host shifts as alternative modes of evolution.

### Reannotation of mildew genomes provides family-level insight into effector gene expansion and loss of core ascomycete genes

Direct cross-species comparisons of gene content and effector repertoires across 50 re-annotated Erysiphaceae genomes revealed patterns of effector diversification and gene family evolution throughout the Erysiphaceae.

The arsenal of effectors available to powdery mildew species has diverged considerably between mildew genera, reflecting lineage-specific evolutionary trajectories within the Erysiphaceae. As reported previously (28,31), our reannotation of the *Blumeria* spp. predicted extremely large effector complements of 452 - 653 genes (154 - 263 EffectorP CSEPs; 350 - 503 homologs to validated effectors) (Figure 2; S7). In contrast, reannotated *Golovinomyces* assemblies carried only 322 - 350 genes (60 - 76 EffectorP CSEPs; 270 - 288 validated effector homologs), consistent with earlier observations of a reduced effector repertoire in dicot-infecting lineages (Wu et al., 2018).

**Fig. 2.**
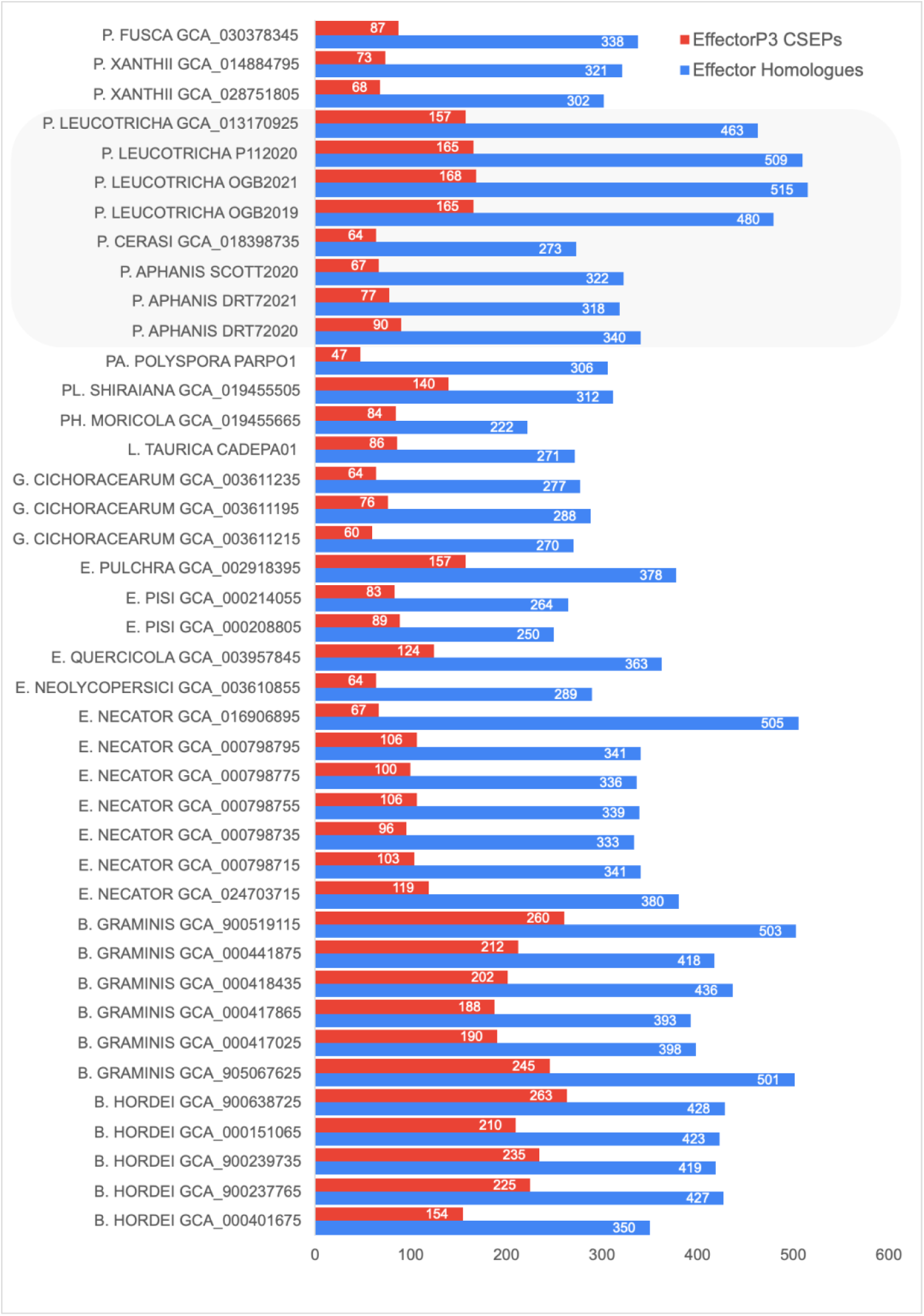
Effector complements across powdery mildew pathogens: Number of Candidate Secreted Effector Proteins (CSEPs) with EffectorP3 support (red) and number of validated effector homologues, including RALPH and EKA homologues (blue) are given based upon *ab initio* gene annotations of powdery mildew pathogen genomes from the *Blumeria, Golovinomyces, Erysiphe, Podosphaera, Leveillula, Pleochaeta* and *Phyllactinia* genera.

Whilst *Blumeria* and *Golovinomyces* species had similar within-genus effector profiles*, Podosphaera* genomes exhibited considerable variation in effector complements, even between closely related species. *P. aphanis* genomes contained comparatively few effectors, with 67 - 90 EffectorP CSEPs and 318 - 340 validated effector homologues for a total effectorome size of 362 - 392 genes (similar to *Golovinomyces* and *Erysiphe* genera). In contrast, *P. leucotricha* assemblies carried 157 - 168 EffectorP3 supported CSEPs and 463 - 515 validated effector homologues, for a total effectorome of 532 - 576 genes (Figure 2; S7). Thus, *P. leucotricha* harbours more than twice as many CSEPs, ∼50% more effector homologues, and ∼50% more total effectors than *P. aphanis.* The average effectorome size across the three *P. leucotricha* assemblies (561 genes) was similar to that of *Blumeria* spp. (585 genes).

These observations indicate that effector expansion in powdery mildew pathogens cannot be attributed solely to broad host type (e.g. monocot versus dicot) or taxonomic position but also reflects lineage-specific evolutionary pressures. The *Podosphaera* genus therefore exemplifies how even closely related biotrophic fungi can develop along distinct evolutionary trajectories.

Alongside effector expansion, a shift to parasitism in phytopathogens is often associated with the loss of superfluous genes no longer required after the transition to host-obligate parasitism. Assessment of gene family size across the powdery mildew clade revealed gene losses predominate early in Erysiphaceae evolution (Figure 3). For example, 527 gene families contracted following the separation of the early diverging species *Pa. polyspora,* while only two showed evidence of expansion. Contractions also predominate at the division of each mildew genera, eg. *Blumeria* (-646)*, Podosphaera* (-405), and *Erysiphe* (-99). These results suggest that the switch to obligate biotrophy in the Erysiphaceae has precipitated gradual ongoing gene losses, whilst species-specific expansions likely reflect subsequent host-specific adaptations.

**Fig. 3.**
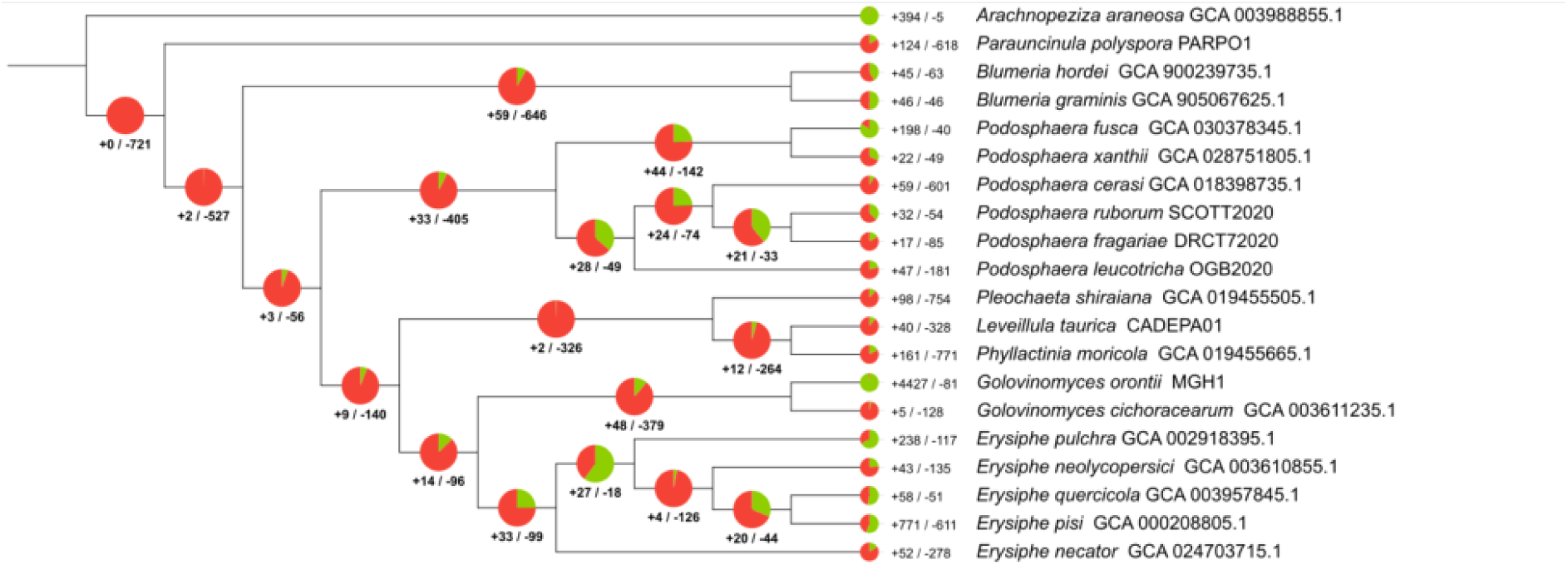
Divergence of powdery mildew pathogens is associated with reductions in gene family sizes: Phylogeny of genome-sequenced isolates from the Erysiphoid clade, including Erysiphaceae species (powdery mildew pathogens) and *Arachnopeziza araneosa* (saprobe). The number of gene families predicted by orthofinder to have increased (green) and decreased in size (red) are shown at each node of the species phylogeny.

A set of 99 purportedly Missing Conserved Ascomycete Genes (MCAGs) has previously been defined for powdery mildew pathogens (Spanu et al., 2010). Our analysis of these 99 MCAGs across the mildew clade and closely related species from the class Leotiomycetes confirmed that many conserved genes have been lost in Erysiphaceae species whilst most MCAGs are present in other fungal families (S8). Additionally, and consistent with results reported by Spanu et al. (2010), a subset of MCAGs were identified as missing from non-mildew biotrophic *Puccinia* species, which share features of powdery mildew pathogen epidemiology, i.e. haustorium formation. Reimplementing the established one-way BLAST-based methodology for MCAG identification suggested discernible differences in patterns of gene loss between mildew genera (S8). However, reciprocal BLAST analysis indicated that many hits represented only distant or domain-level similarities rather than bona fide homologues. Nonetheless, the relative retention of certain ancestral genes in *Podosphaera* supports the view that genome reduction during the evolution of obligate biotrophy proceeds in a lineage-specific and stepwise fashion.

### Evidence-guided gene prediction provides insight into species-specific effector gene expansion in *Podosphaera leucotricha*

RNA-seq data were utilised for a *de novo* annotation of *P. aphanis* and *P. leucotricha* assemblies (Table 2). Predicted gene counts in *P. leucotricha* assemblies ranged from 17,993 - 18,651, whilst for *P. aphanis*, 14,829 - 17,239 genes were predicted from strawberry samples, while the raspberry sample SCOTT2020 contained 15,056 genes. Similarity and HMM searches identified *P. leucotricha* and *P. aphanis* proteins homologous to effectors in PHI-base or the curated Predector database, primarily from *B. graminis* and *B. hordei*. Among these were 156–165 non-canonically secreted EKA effector homologues in *P. leucotricha* and 72 - 103 in *P. aphanis* (Table 2), consistent with lineage-specific effector diversification between even closely related *Podosphaera* genera.

**Table 2.**
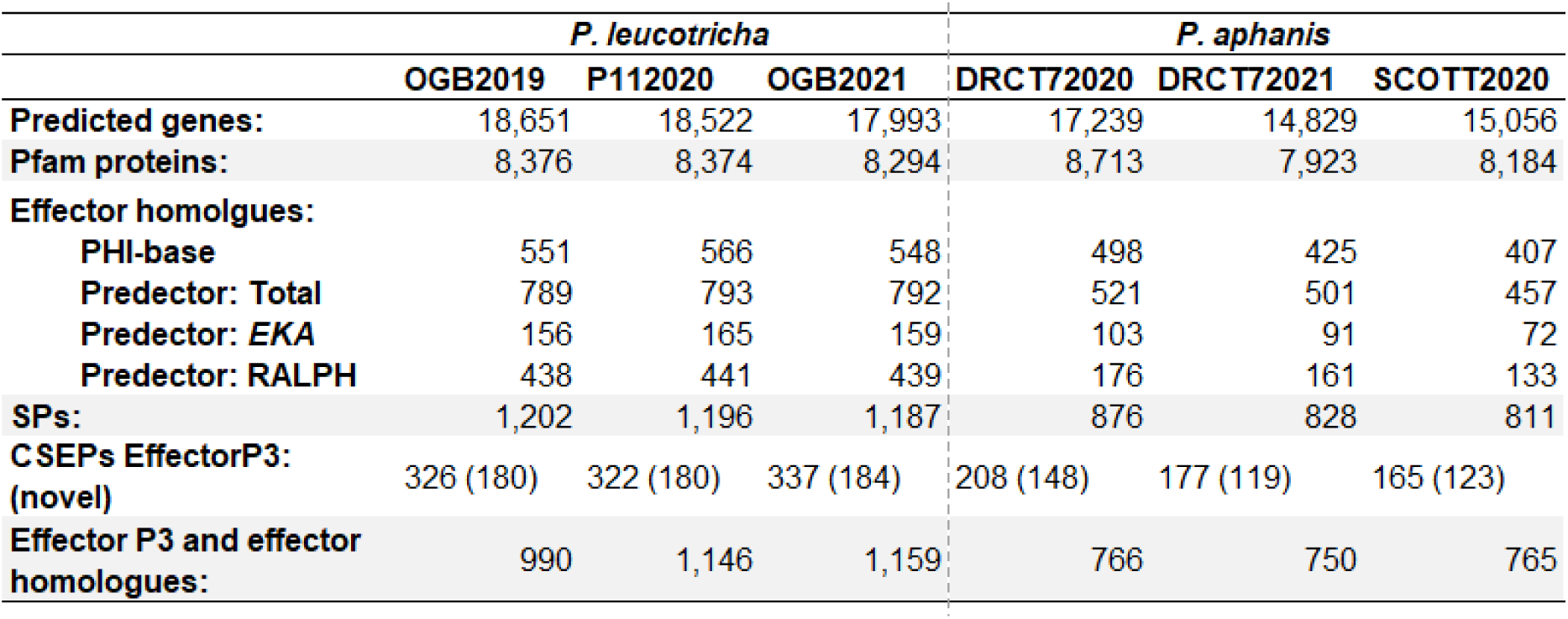
*Podosphaera aphanis* and *Podosphaera leucotricha* proteome annotations: Total gene models and number of functionally annotated genes by effector-family across assembled *Podosphaera* genomes are given. Including: the number of gene predictions; the number of resulting proteins with annotations in the Pfam database; with homology to pathogenicity-related genes in PHI-base; and to a database of previously-characterised effector genes supplied in the Predector pipeline including both EKA and RALPH homologues. Predicted numbers of Secreted Proteins (SPs) and Candidate Secreted Effector Proteins (CSEP) with EffectorP3 predictions of effectors are given, and in brackets, the number of these proteins which have no hits in PHI-base or the collection of known effector proteins. Finally the number of putative effectors from all annotation methods.

|  | *P. leucotricha* |  |  | *P. aphanis* |  |  |
| --- | --- | --- | --- | --- | --- | --- |
|  | OGB2019 | P112020 | OGB2021 | DRCT72020 | DRCT72021 | SCOTT2020 |
| **Predicted genes:** | 18,651 | 18,522 | 17,993 | 17,239 | 14,829 | 15,056 |
| **Pfam proteins:** | 8,376 | 8,374 | 8,294 | 8,713 | 7,923 | 8,184 |
| **Effector homologues:** |  |  |  |  |  |  |
| **PHI-base** | 551 | 566 | 548 | 498 | 425 | 407 |
| **Predector: Total** | 789 | 793 | 792 | 521 | 501 | 457 |
| **Predector: EKA** | 156 | 165 | 159 | 103 | 91 | 72 |
| **Predector: RALPH** | 438 | 441 | 439 | 176 | 161 | 133 |
| **SPs:** | 1,202 | 1,196 | 1,187 | 876 | 828 | 811 |
| **CSEPs EffectorP3:**<br>**(novel)** | 326 (180) | 322 (180) | 337 (184) | 208 (148) | 177 (119) | 165 (123) |
| **Effector P3 and effector**<br>**homologues:** | 990 | 1,146 | 1,159 | 766 | 750 | 765 |

Orthology analysis of predicted proteins from the six *de novo* assemblies identified conserved and species-exclusive gene families. Orthogroups shared among all three assemblies of a species were considered species-conserved, with *P. leucotricha* assemblies demonstrating a significantly greater proportion of species-conserved orthogroups (86.7% (95% CI 86.2 - 87.2%)) than *P. aphanis* (69.6% (95% CI 68.8 - 70.4%)) (Figure 4). Within *P. aphanis*, divergence between strawberry and raspberry isolates was evident, with 15.3% of orthogroups shared between the two strawberry assemblies but absent from the raspberry assembly (Figure 4A), confirming host-associated differentiation within the *P. aphanis* complex. Taking the DRCT72020 strawberry (*P. aphanis*) and P112020 apple (*P. leucotricha*) assemblies as species representatives, 2,147 *P. aphanis*-exclusive and 4,702 *P. leucotricha*-exclusive gene predictions were identified, underscoring substantial lineage-specific gene innovation within *P. leucotricha*.

**Fig. 4.**
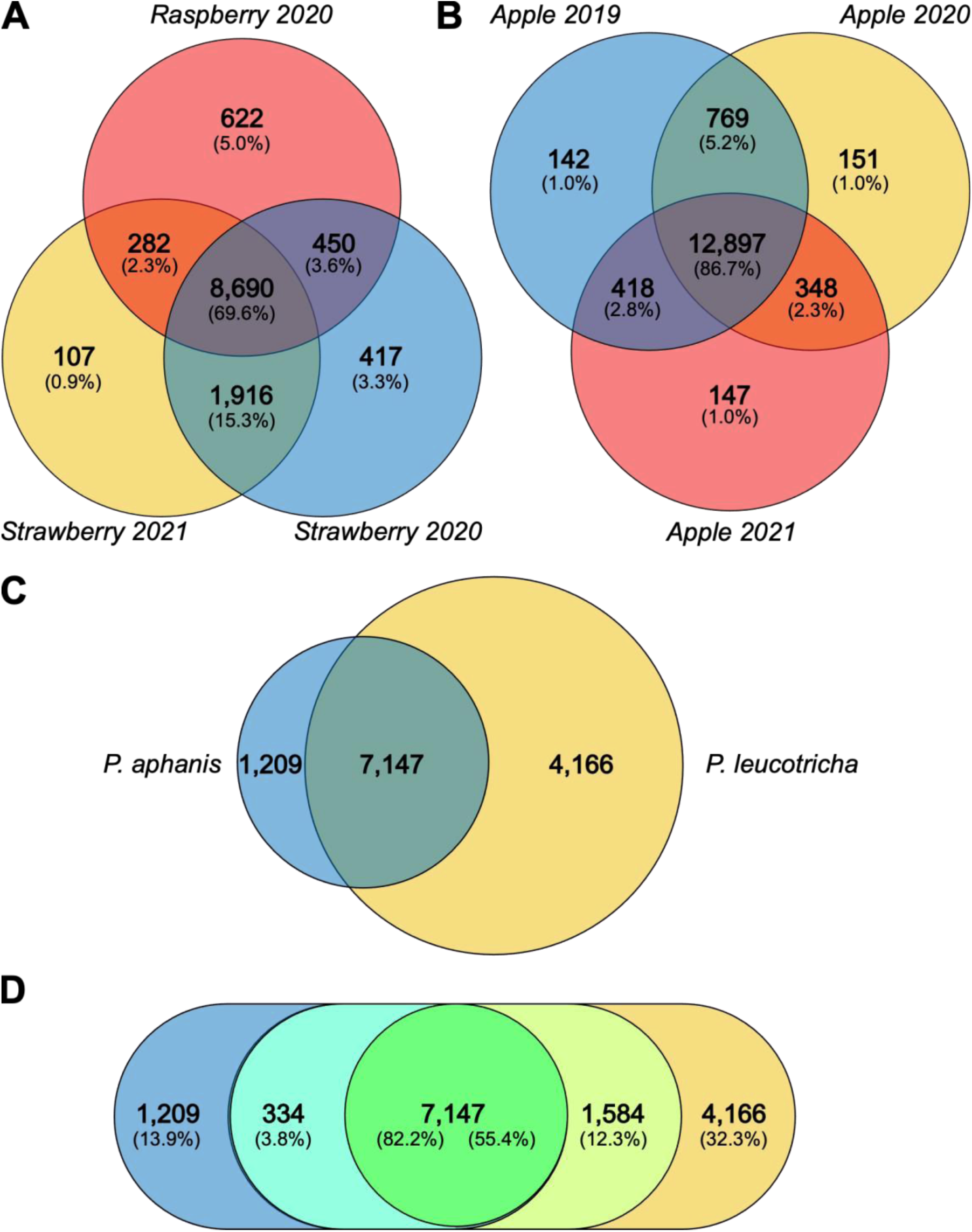
Podosphaera *aphanis* and *Podosphaera leucotricha* unique and common orthogroups: Venn and Euler diagrams presenting the presence/absence of orthologous gene groups across the genomes : the three *P. aphanis* assemblies (A); the three *P. leucotricha* assemblies (B); orthogroups unique to *P. aphanis* or *P. leucotricha* assemblies, or found in both species (C); orthogroups unique to *P. aphanis*/*P. leucotricha* assemblies or common to all six assemblies (D).

Among the candidate *P. aphanis*–exclusive genes, 43 encoded proteins matched experimentally validated fungal effector HMM profiles, whereas five times more (210) *P. leucotricha*–exclusive effector homologues were identified (S9). Similarly, *de novo* effector prediction identified only 38 CSEPs within *P. aphanis*-exclusive orthogroups, but over a hundred in *P. leucotricha*. This disparity was not attributable to divergence between strawberry and raspberry isolates, as excluding the SCOTT2020 raspberry assembly only increased *P. aphanis* CSEP count to 62 (S10). Together, these data show uneven effector diversification between *Podosphaera* species, suggesting that *P. leucotricha* has undergone more extensive effector innovation than *P. aphanis*.

*ssssP. aphanis-* and *P. leucotricha-*exclusive CSEPs were searched against genomes from the wider Erisiphaceae in order to validate their designation as species-specific effectors (Figures 5 and 6). Ten *P. aphanis*-exclusive CSEP genes were confirmed as specific to *P. aphanis*, whilst the remaining 28 genes highlighted by orthology analysis displayed restricted distributions within the *Podosphaera* clade and, notably, *L. taurica*, which causes powdery mildew on sweet pepper (Figure 5). Not all *P. aphanis-*exclusive CSEPs were present at low-copy number when searched against assemblies, with several returning >10 BLAST hits in *P. aphanis*, *L. taurica*, *P. xanthii*, and *P. fusca* assemblies.

**Fig. 5.**
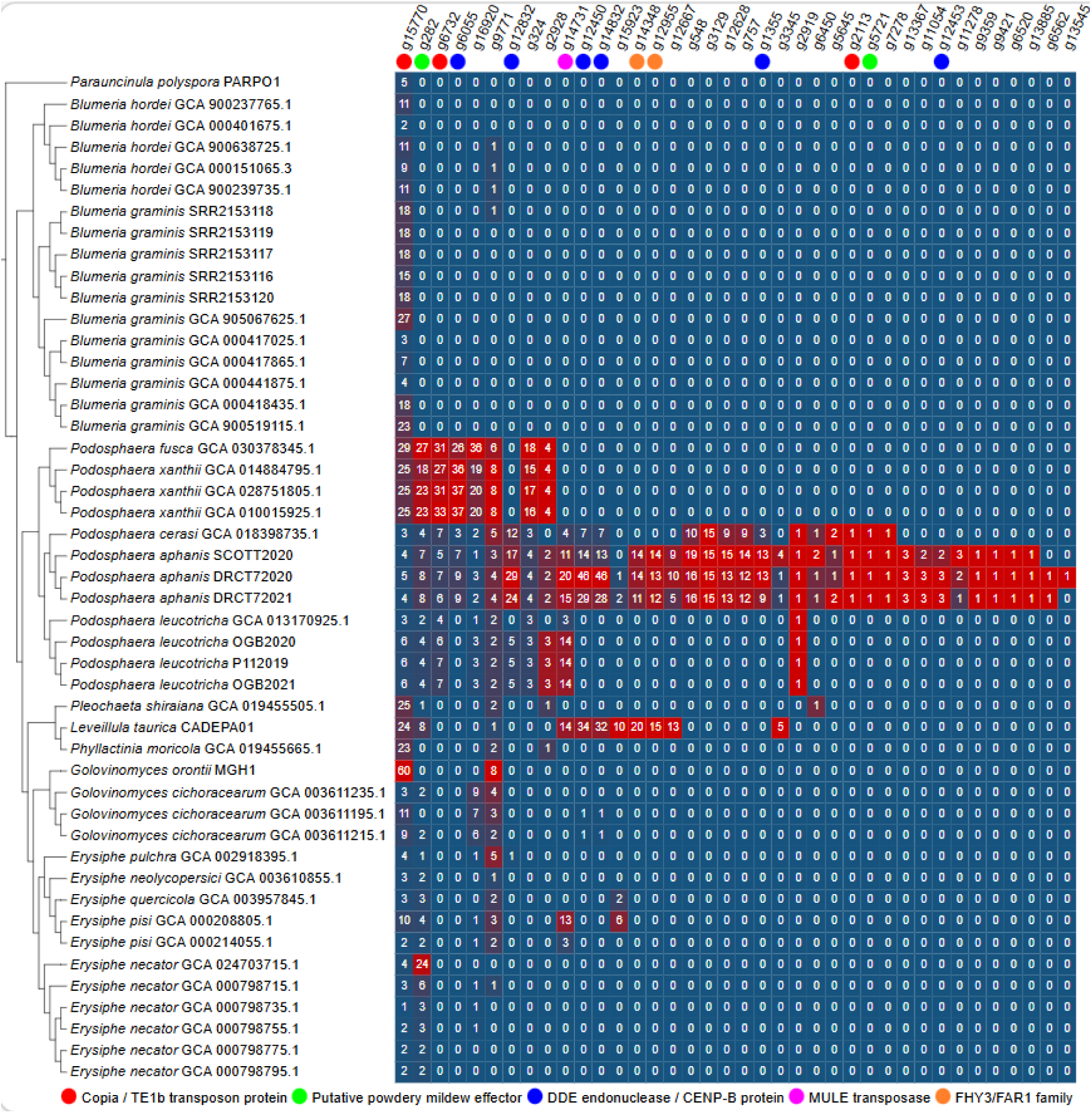
Confirmation of species-specific effector candidates through TBLASTN analysis: A phylogeny of Erysiphaceae (powdery mildew pathogen) assemblies is presented, for each given *Podosphaera aphanis* EffectorP3 supported Candidate Secreted Effector Protein (CSEP) the number of TBLASTN hits in each available Erysiphaceae genome (S2) is given, shading is normalised for each gene to highlight increased occurrence. BLASTP hits including alignments to putative effectors in other powdery mildew species and transposon associated/derived proteins are indicated.

**Fig. 6.**
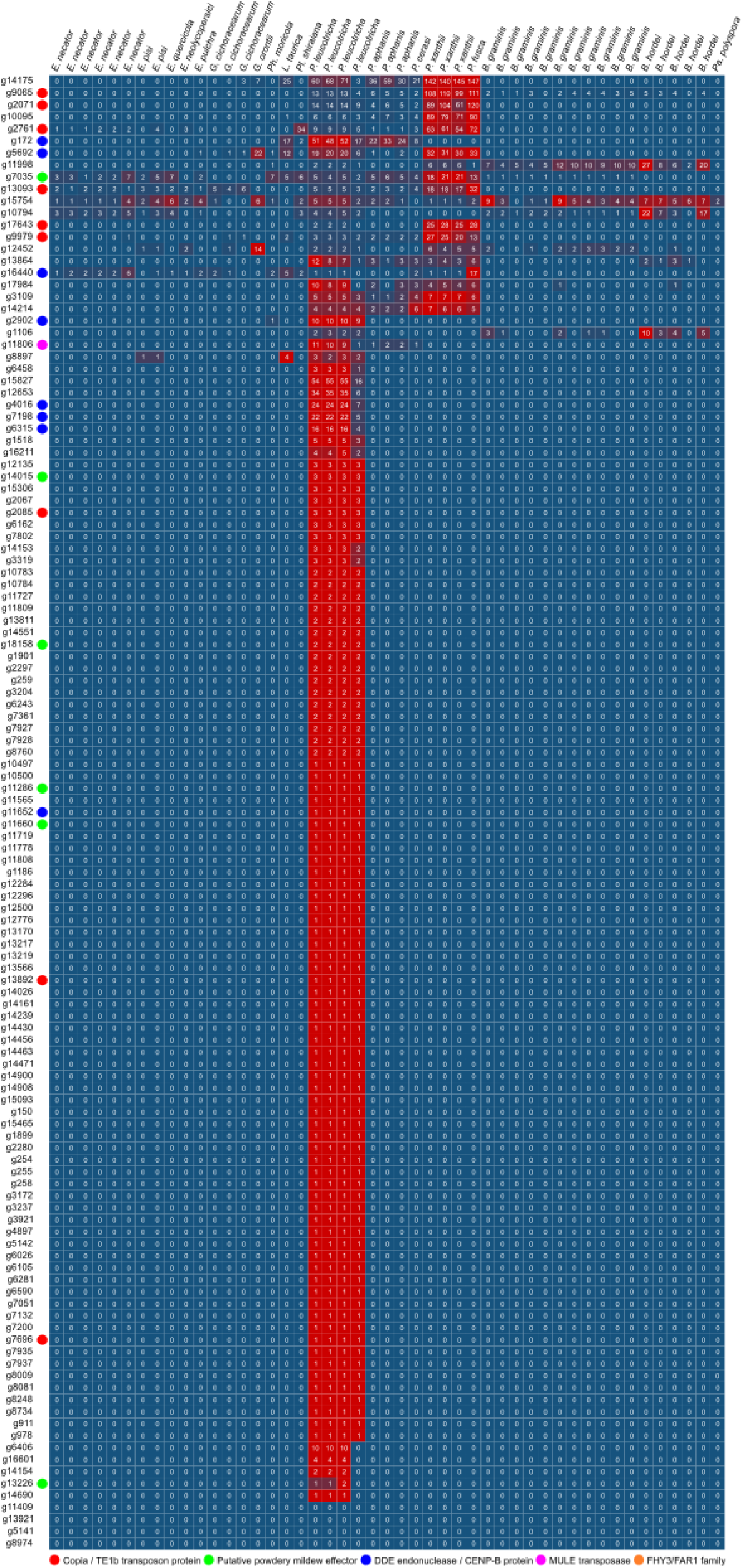
Confirmation of species-specific effector candidates through TBLASTN analysis: For each given *Podosphaera leucotricha* EffectorP3 supported Candidate Secreted Effector Protein (CSEP) the number of TBLASTN hits in available Erysiphaceae (powdery mildew pathogen) assemblies (S2) is given, shading is normalised for each gene to highlight increased occurrence. BLASTP hits including alignments to putative effectors in other powdery mildew species and transposon associated/derived proteins are indicated.

In contrast to *P. aphanis*, *P. leucotricha* carried many species-specific CSEPs occurring in low (<3) copy number (Figure 6). Of 123 EffectorP3 supported CSEPs predicted from *P. leucotricha*-exclusive orthogroups 57 had one hit, and 16 two hits, in all *P. leucotricha* assemblies but no other species, and a further five had hits in all in-house *P. leucotricha* assemblies only. As in *P. aphanis,* a subset (18) of *P. leucotricha*-exclusive effector candidates occurred at elevated copy number in one or more powdery mildew genomes.

The majority of species-specific CSEP lacked annotations; however, GO and EGGNOG terms related to ribonuclease activity and nucleic acid binding were common where annotations were present. An RNase A-like fold is a conserved feature of the RALPH effector family, and all proteins with putative ribonuclease domains were also identified as RALPH homologues. These proteins likely represent novel species-specific RALPH variants, consistent with continued diversification of this effector family across powdery mildew fungi. BLAST searches against NCBI nr database returned low percentage identity alignments to ribonuclease and putative effectors in other powdery mildew species for some CSEPs, other CSEPs returned alignments to transposon associated and derived proteins (eg. DDE superfamily endonuclease, GAG-pre-integrase, MULE transposase, centromere protein B, FHY3/FAR1 family). These transposon related alignments were concentrated amongst CSEPs with elevated copy number (>10 BLAST hits) (Figure 6), suggesting that whist they fulfil the criteria of CSEPs these genes may represent mobile element fragments rather than functional effector genes.

## DISCUSSION

The evolution of obligate biotrophy is characterised by both the expansion of lineage-specific effector repertoires and the loss of otherwise conserved genes (31,62,63). These complementary processes shape how pathogens interact with and overcome host defences. Effector expansion has been considered a hallmark of monocot-infecting powdery mildew species, whereas dicot-infecting powdery mildew pathogens were thought to carry smaller effector repertoires (32). However, this view is not supported by our comparative analysis of two closely related *Podosphaera* species. We showed that *P. leucotricha*, a host-specific dicot pathogen of apple, possesses an effectorome comparable in size to those of monocot powdery mildew pathogens, including expanded families of RALPH and non-canonically secreted EKA effectors. In contrast, *P. aphanis,* which infects strawberry and raspberry, had a much smaller effectorome. These results demonstrate that effector expansion amongst the Erysiphaceae cannot be predicted solely by host type (monocot vs. dicot), host range, or genus, but instead reflects more complex and species-specific evolutionary pressures. Additionally, we highlighted effector candidates unique to *P. aphanis* and *P. leucotricha*, which likely confer their distinct host specificities and present good targets for diagnostic testing. In parallel, we observed the loss of many genes conserved in non-mildew fungi, and the stepwise contraction of gene families between powdery mildew genera. These ongoing losses likely reflect continuing adaptation to obligate parasitism of particular hosts, which renders many genes redundant or potentially deleterious if detectable by the host immune system. Taken together our findings highlight contrasting effects of biotrophic adaptation on pathogen genomes: diversification of effectoromes in response to host immune landscapes, and the convergent loss of conserved fungal genes.

### Annotation of fifty mildew genomes provides family-level insight into effector gene expansion

Both *P. aphanis* and *P. leucotricha* possess larger effector repertoires than reported previously for dicot powdery mildew pathogens (29,32). However, we also observed a large difference in effectorome size between *P. leucotricha* and *P. aphanis,* despite both species exhibiting narrow host ranges and being closely related. The proliferation of CSEPs in *Blumeria* spp. has been attributed to the presence of allelic series of *R* genes in wheat and barley (64). A similar mechanism likely underlies effectorome expansion in *Podosphaera*. Numerous major powdery mildew *R* genes have been described in apple (Lesemann, 2011), in contrast, only quantitative resistance, varying with season and clonal propagation, is known in strawberry (65,66). Consistent with this, a complex race structure has been reported in apple *P. leucotricha*, where isolates vary in their ability to infect different apple cultivars (67,68), whereas evidence for races in *P. aphanis* is limited to resistance breaking in the cultivar ‘Korona’ (69). Therefore, effectorome expansion observed in *P. leucotricha* may reflect stronger host-driven selective pressures within the apple pathosystem.

### RALPH and EKA effectors are prevalent across *Podosphaera*, with recent expansion in *Podosphaera leucotricha*

*RALPH* genes are a family of effectors common across powdery mildew species (27,70). The gene family shares a common RNAse fold and is thought to have descended from a single ancestral gene encoding a ribonuclease. The presence of numerous RALPH orthologues in the predicted *proteomes of P. aphanis and P. leucotricha* confirms that this progenitor ribonuclease was a relatively ancient gene. Previous work predicted a similar number of RNase-like proteins in the proteomes of monocot-infecting *Blumeria* spp. and dicot-infecting powdery mildew pathogens (32). However, the secretomes of *Blumeria* spp. displayed a pronounced enrichment in RNase and hydrolase activities relative to dicot-infecting species, leading to far more RNase-like genes being designated as RALPH CSEPs in monocot-infecting *Blumeria* spp. (32).

In this study, we performed homology-based and feature based (CSEP) annotation of putative effectors in parallel. Homology based annotation found that the *RALPH* gene family was expanded in *P. leucotricha.* The *P. leucotricha* genomes encoded triple the number of RALPH homologues found in strawberry infecting *P. aphanis*, and raspberry infecting *P. aphanis* appears to carry an even smaller set (Table 2). In addition, many (240) *P. leucotricha*-specific CSEPs are RALPH homologues (S10). The expansion of the *P. leucotricha-*specific effectorome is therefore reasonably driven by an expansion in the *RALPH* family.

*EKA* genes encode a distinctive powdery mildew effector family that lack N-terminal signal peptide sequences and are therefore presumed to be secreted via ‘non-conventional’ pathways (71,72). As such, *EKA*s escape standard CSEP-prediction pipelines and require homology- or HMM-based searches for reliable detection (71–74). Most *Blumeria* effector gene families are thought to have arisen after the divergence of *Blumeria* from other powdery mildew clades ∼76 million years ago (73,75). However, we identified EKA homologues in both *P. leucotricha* and *P. aphanis* proteomes (Table 2), suggesting that the *EKA* family originated prior to the divergence of *Blumeria* (11). The *EKA* family appears to be expanded in the *Podosphaera* genus, with >150 homologous proteins predicted from the *P. leucotricha* genome and >70 homologues predicted from *P. aphanis* (Table 2). *EKA* genes are thought to have evolved from 3’-truncated ORF1 proteins of class I long-interspersed element retrotransposons (76). Notably, LINE retrotransposons make up a similar proportion of *B. hordei* (6.5%) (76) and *P. leucotricha* & *P. aphanis* assemblies (5 - 6%), this may help to explain the large number of *EKA* homologues observed in these species.

### Effector evolution in powdery mildew pathogens

High copy number CSEP candidates may in fact represent transposon sequences. In support of this hypothesis, in addition to fulfilling CSEP criteria (secretion signal and no homology in unrelated species), annotations for transposon-related domains were also common among high copy number candidates. Lineage-specific transposons may form a stepping stone towards novel lineage-specific virulence factors in powdery mildew pathogens. Several powdery mildew species display evidence of recent or ongoing TE activity (21,61,64). TEs are extremely lineage-specific and likely contribute to shaping the genomes of different powdery mildew species in a lineage-specific manner (77). A ‘devil’s bargain’ has been described between TEs and plant pathogens (78). Uncontrolled TE proliferation leads to genome instability. On the other hand, TEs are recognised as drivers of mutational variation that fuel adaptation, and are closely associated with effectors in fungal plant pathogens including powdery mildew pathogens (79). TE insertions may lead to increased virulence by disrupting avirulence genes but have also been connected to the evolution of novel virulence factors (80,81). TEs can contribute to the evolution of effector genes by donating novel lineage-specific regulatory elements to preexisting host genes or by losing their ability to self-replicate and transitioning to become novel host genes through a process called molecular domestication (77,82). The association of transposon related domains with a secretion signal could represent an important step in the development of novel virulence factors.

EKA and ROPIP1 powdery mildew effector genes are thought to have evolved from TEs in *B. hordei* (76,83). ROPIP1 originates from a SINE/Eg-R1 TE that is exceptionally common in the genome of *B. hordei* and interacts with susceptibility factor RACB in barley host cells to support haustorium establishment (83). Whereas it is hypothesised that *EKA* genes are derived from the ORF1 gene of a LINE retrotransposon (72,76,84). However, none of the transposon annotated genes in *P. aphanis* or *P. leucotricha* are homologous to known validated effectors such as the *EKA* family.

### Gene losses across the powdery mildew pathogens

Our analyses highlight widespread gene losses accompanying both the initial shift to obligate parasitism and on an ongoing basis in different mildew genera, supporting extensive gene family contraction as a defining feature of Erysiphaceae genome diversification. This trend is consistent with previous reports of gene loss in other obligate biotrophs, where genes dispensable in a host-dependent lifestyle are gradually eliminated (31,57,85,86). While gene family contractions were dominant at higher taxonomic levels, species-specific expansions were apparent, including in *P. aphanis, B. hordei,* and *B. graminis.* These expansions likely reflect species-specific adaptations, possibly associated with host specialisation and the fine-tuning of parasitic strategies, as seen in the expansion of putative effectors. The contrast between broad-scale contraction of gene families but lineage-specific expansion of some groups demonstrates that the evolution of obligate biotrophs is potentially driven by at least two complementary processes: the loss of both deleterious and superfluous genes, and the gain of new genes conferring advantages in a parasitic lifestyle.

## CONCLUSIONS

Analysis of 50 Erysiphaceae genomes revealed substantial gene gains and losses linked to lineage divergence and host–pathogen coevolution in the Erysiphaceae. Focused analyses of six *P. aphanis* and *P. leucotricha* assemblies revealed expansion of RALPH and EKA effector families. There were striking differences between these two closely related species with effector expansion far more pronounced in *P. leucotricha.* We also confirm the widespread loss of conserved ascomycete genes across the mildew clade, likely reflecting progressive genomic streamlining associated with the evolution of obligate parasitism, together with ongoing gene family contraction in multiple lineages. By characterising the effector repertoires of apple and strawberry mildews, this study provides new insight into how genomic innovation and reduction interact to shape the evolution of obligate biotrophs. The resulting genome resources and effector catalogues will accelerate the discovery and functional validation of host resistance genes, ultimately supporting breeding strategies for durable powdery mildew resistance in apple and strawberry.

## DATA AVAILABILITY

This Whole Genome Shotgun project has been deposited at DDBJ/ENA/GenBank under the accessions JALNPU000000000, JALMLP000000000, JALMLQ000000000, JAMDJL000000000, JBVYIF000000000 an JAKRRZ000000000. The version described in this paper is version XXXXXX010000000 for JBVYIF010000000 and versions XXXXXX020000000 for all additional genomes. Illumina sequence data associated with this work has been deposited on the Sequence Read Archive (SRA) under BioProject PRJNA744412.

## ACKNOWLEDGEMENTS

Members of the pathology team at NIAB East Malling site are recognised for support in field and pathology work covered in this manuscript. Whereas members of the Armitage Lab, Cockerton Lab are recognised for critical project discussion.

## S1: SUPPLEMENTAL METHODS

### DNA Extraction

High-molecular-weight DNA extractions were performed based on the CTAB extraction protocol of Schwessinger (2016), which was modified to obtain high-quality DNA from mildew. Premade buffers were combined to form lysis buffer (Buffer A: 0.35 M sorbitol; 0.1 M TrisHCl; 5 mM EDTA pH 8 ● Buffer B: 0.2 M Tris-HCl; 50 mM EDTA pH 8; 2 M NaCl; 2% CTAB ● Buffer C: 5% Sarkosyl N-lauroylsarcosine sodium salt ● Buffer D: PVP40 10% ● Buffer E: PVP10 10%) in the ratios 5:5:2:1:1, 10 µL (10 kU) Rnase A was added per 14 ml of lysis buffer (1). Fungal samples were combined with silicon dioxide sand, 50 - 70 mesh particle size, (1:1) and ground under liquid nitrogen using a mortar and pestle. Samples were then allowed to equilibrate to room temperature before being combined with lysis buffer in a 1:8 ratio. Samples were then incubated at room temperature for 30 min whilst being inverted, 15 µL of proteinase K was then added per ml of lysis suspension and room temperature incubation continued for another 30 min. Following this, samples were cooled on ice for 5 min before 5 M potassium acetate was added in a 1:5 ratio and cooling continued for an additional 5 min. Samples were then centrifuged for 12 min at 5,000 × g, and the supernatant was collected. Washing was then carried out by the 1:1 addition of Phenol:Chloroform:Isoamyalcohol 100 mM Tris-EDTA pH 8 (P:C:I). Samples were mixed by inversion for 2 hours and centrifuged for 10 min at 4,000 × g before transferring the supernatant to a fresh tube. This wash was repeated twice per sample, followed by a third wash using Chloroform:Isoamyl alcohol (C:I) in place of P:C:I. DNA precipitation was performed by adding 1:10 Sodium Acetate (3 M pH 5.2) and 1:1 Isopropanol, followed by overnight incubation at 4 °C. DNA was then pelleted by centrifugation at 8,000 × g for 30 min. The supernatant was discarded, and the pellet washed three times by resuspension in 1.5 ml of 70% ethanol, centrifugation at 13,000 × g for 5 min, and discarding of the supernatant. Following the final wash step, the remaining ethanol was allowed to evaporate for 30 min before the DNA pellet was dissolved in 200 µL of 10 mM Tris pH 8.5 at room temperature for three hours. The quality of extracted DNA was initially assessed using a Nanodrop 1000 spectrophotometer (Thermo Scientific), provided these results were in the target range (OD260/230 = 1.8 - 2.0, OD260/280 = 1.8 - 2.0) then quantity and RNA contamination were further assessed via Qubit dsDNA High-sensitivity and Qubit RNA High-sensitivity assay kits with a Qubit 3.0 fluorometer (Life Technologies, Waltham, MA USA).

### ITS Sequencing

The ITS regions of isolates were amplified using serial dilutions of DNA samples. PCR was performed using 5 µL of Taq 5X master Mix (NEB), 16 µL water, 1 µL ITS-1 primer (TCCGTAGGTGAACCTGCGG), 1 µL ITS-4 primer (TCCTCCGCTTATTGATATGC) and 2 µL DNA dilutions. PCR was performed on a Veriti thermal cycler (Applied Biosystems) using the following cycling conditions: initial 95 °C for 3 min; 35 cycles of 95 °C for 20 s, 60 °C for 15 s, and 68 °C for 2 min; and final extension at 68 °C for 2 min. PCR products were visualised on a 1.5% agarose gel with GelRed (0.5 µL L-1) before being purified using a Monarch® PCR & DNA clean-up kit (5 µg) following the manufacturer’s instructions. Purified PCR products were quantified, diluted, and combined with forward and reverse primers in separate tubes before submission for Sanger sequencing via the Eurofins Genomics LIGHTrun tube service. The resulting sequences were aligned to a reference ITS region downloaded from NCBI (GenBank accession no.: JQ999954 for P. leucotricha and KT359262 for P. aphanis) and analysed using Geneious V10.0.2.

### RNA Extraction

RNA was extracted for RNA-seq and gene annotation. RNA extractions were performed using 3% CTAB extraction buffer as described by Yu et al. (2012) with the following modifications: chloroform:isoamyl alcohol (24:1) washing was omitted; and precipitation was performed at -20 °C for four hours (2). The resulting RNA concentration and RNA Integrity Number (RIN) of samples were assessed using the Agilent RNA ScreenTape System with a 2,200 Tapestation (Agilent Technologies, Germany) according to the manufacturer’s protocols. DNA contamination was assessed via Qubit dsDNA HS assay kit with a Qubit 3.0 fluorometer (Life Technologies, Waltham, MA, USA).

### Genome Assembly

The mildew genome assembly pipeline used was as follows. Raw sequencing reads were trimmed, and adapters removed using Trimmomatic v0.39 (3). Reads were subjected to a quality control check using FastQC v0.11.9, before and after trimming (4). Kraken 2 v2.1.1 was used to taxonomically classify these trimmed reads (k-mer length = 31) (5). To do this, a custom database was constructed, including the standard Kraken 2 databases for archaea, bacteria, fungi, plants, protozoa, viruses, and vertebrate mammals, which included the apple genome, with the addition of other host plant genomes, potential contaminants, and 29 Erysiphaceae genomes publicly available in the NCBI database. Taxonomic classifications were visualised using Pavian v1.0 (6). Coverage was estimated using SAMtools v1.1 ‘coverage’ function and the K-mer Analysis Toolkit v2.4.2 function ‘kat plot spectra-cn’ (7,8). Following this, reads were aligned to the respective host genome for each sample: apple, strawberry, or raspberry (9–11).

Those reads not aligning to the host genome were carried forward for de novo genome assembly using SPAdes v3.14.1 (12). The custom Kraken 2 database was then used to taxonomically classify contigs post assembly (visualised by Pavian) (5,6). All contigs that were not assigned to the class Leotiomycetes were removed. Quality of the resulting genomes was assessed by searching for Benchmarking Universal Single-Copy Orthologues (BUSCO) with BUSCO v4.0.6 (13). For size estimation, 21-mers were counted using Jellyfish v2.2.3 (14). Final assemblies were assessed for contamination using BlobTools: raw sequencing reads were Bowtie 2 aligned to final assemblies to generate coverage files; hit files were generated via a BLASTN search of assembled contigs against the NCBI ‘nt’ database (15). BLASTN classifications were performed to the phylum level, with the Erysiphaceae family belonging to the phylum Ascomycota.

### Transposon analysis

De novo prediction of repetitive elements was performed on the assembled contigs. Repeat masking was performed using RepeatModeler v2.0.2 and TransposonPSI (16,17). The resulting annotations of repetitive sequences (GFF format) were used to mask assembly files using BEDTools v2.30.0 (18). In addition, repetitive element families were quantified and kimura distance and repeat category plots via EarlGreyTE v4.0.8, utilising the Dfam 3.7 open database of transposable elements and repetitive DNA families, with RepeatMasker search term “erysiphales” (19).

### Gene Prediction

RNA-seq was performed in order to provide evidence for gene prediction and transcriptome generation. Sequence reads from RNA-seq were subjected to a quality control check using FastQC v0.11.9. Sequences were trimmed and adapters removed using Trimmomatic v0.39. Reads were then aligned to the draft genome assembly using STAR v2.7.3 (20). The BRAKER v1.9 pipeline was then used to make gene predictions using these alignments, and these predictions were supplemented by predictions made using CodingQuarry v2.0 in pathogen mode (21,22). BRAKER gene models were used preferentially, adding CodingQuarry genes present in intergenic regions, as described by Armitage et al. (2018) (23).

### Intergenic distances

To assess compartmentalisation of the *P. leucotricha* and *P. aphanis* genomes, the distance between genes and repetitive elements was investigated. The 5-prime upstream distance (bp) and 3-prime downstream distance to the nearest gene was counted for each gene model in each assembly. Intergenic distances were plotted for all genes using the R package ggplot2 v3.5.1, additionally BUSCO, CAZY, CSEP (EffectorP3 supported), EKA, and RALPH genes were highlighted (24). Where genes had both 5’ and 3’ measurements total intergenic distances were extracted via a custom python script (https://github.com/TCHeaven/Scripts/tree/main/gruffalo/find_intergenic_regions.py). Five prime and three prime intergenic distances were inspected for BUSCO genes, QQ plots and Shapiro-Wilk test demonstrated that intergenic distance residuals were not normally distributed. Therefore, permutation tests were performed in R with 10,000 iterations to compare the intergenic distances of putative CAZY, CSEP, EKA, and RALPH genes to a control group of Leotiomycete BUSCO gene orthologues (25). Due to the fragmented nature of the assemblies many genes were located on the ends of contigs and therefore did not have an up- or down-stream neighbouring gene, these missing distances were assigned a value of 99,999 for plotting purposes and were excluded from permutation testing. The simpler Mann-Whitney U test was used to investigate differences between three and five prime intergenic distances as dataset size and distributions are similar for the two measurements.

### Analysis of Phylogenetic Relationships Between Mildew Taxa

Phylogenies were constructed for powdery mildew pathogen and host species. The resulting trees were visualised using iTOL v5 with connections between pathogens and hosts drawn by hand in inkscape v1.4.2 (26).

Benchmarking universal single-copy orthologues were used to construct phylogenies for mildew assemblies and a selection of other Helotiales spp.. Leotiomycete genes (leotiomycetes_odb10 BUSCO database) were extracted and orthologues were retained if a single hit was found in more than three Erysiphaceae genomes. Nucleotide sequences from the resulting 3,232 Leotiomycetes hits were aligned using MAFFT v7.490, before being trimmed using trimAl v1.4.rev15 (27,28). IQ-TREE v2.3.0 (29) was used to select a best-fit model via the “-m MFP” flag and subsequently to calculate a phylogenetic tree from concatenated BUSCOs with parameters “-m SYM+I+R9 -B 1000”. Additionally, RAxML v8.1.17 was used to calculate the most parsimonious tree for each locus from 1000 bootstraps, a single consensus phylogeny was then determined for pathogen species and for host species using ASTRAL v5.7.8 (30,31). Concatenation-based IQTREE and quartet-based ASTRAL species trees were highly congruent.

In order to generate a phylogeny for subsequent CAFE analysis the IQTREE phylogeny was pruned via iTOL v5 to produce a species tree consisting of 19 Erysiphaceae assemblies rooted to A. araneosa, nodes with only one remaining leaf were collapsed (26). The propose *Podosphaera fragariae* (*P. aphanis* on strawberry) and *Podosphaera roborum* (*P. aphanis* on raspberry) were considered separate species for this analysis. The phylogeny was converted to an ultrametric tree using the penalised likelihood approach implemented by the “chronos” function from the R package ‘ape’. The following time points were used for calibration of divergence time: *B. graminis*-*Pa. polyspora*, 79.0 Million Years Ago (MYA); *B. graminis*-*E. necator*, 53.2 - 75.0 MYA (34–36).

A list of species names was uploaded to TimeTree (http://timetree.org) in order to generate a phylogeny of mildew host plants (37). The TimeTree database could resolve divergence times for all host species except Triticale, the host of B. graminis assembly GCA905067625.1, in addition, substitutions were made for some taxa: *Hevea brasiliensis* (replaced with *Hevea guianensis*); *Celtis sinensis* (replaced with *Celtis occidentalis*); *Morus alba* (replaced with *Morus rubra*); *Fragaria ananassa* (replaced with *Fragaria vesca*); *Pisum sativum* (replaced with *Pisum fulvum*); *Triticum aestivum* (replaced with *Triticum monococcum*).

To place the strawberry and raspberry powdery mildew samples within the *P. aphanis* complex, a phylogeny of the clade was constructed. A representative ITS and 28S sequence (NCBI accession LC777859.1) was queried against in-house *P. aphanis* genome assemblies using BLAST v2.16.0 (38), and matching regions were extracted with Samtools v1.22.1 (7). Reverse-complemented sequences were aligned to the *P. aphanis* dataset of Bradshaw et al. (2025) using MAFFT v7.525 (28,39). IQ-TREE v3.0.1 (29) was used to select the best-fit model with -m MFP, and phylogenetic trees were inferred using the parameters “-m HKY+G4 -B 1000 -alrt 1000”.

## SUPPLEMENTAL MATERIALS, TABLE AND FIGURE LEGENDS

**S1: Supplemental methods** providing additional detail on genomic analyses performed within this manuscript.

**S2: Erysiphaceae genomes included in phylogenies and comparative genomics.** Species name, source of the genome, identifier, and host plant from which the assembly sample was collected. Genomes marked by an asterisk (*) were used in custom kraken2 database construction.

**S3: Non-mildew genomes included in phylogenetics and comparative genomics analyses.** Species name, source of the genome, and identifier are provided.

**S4:**
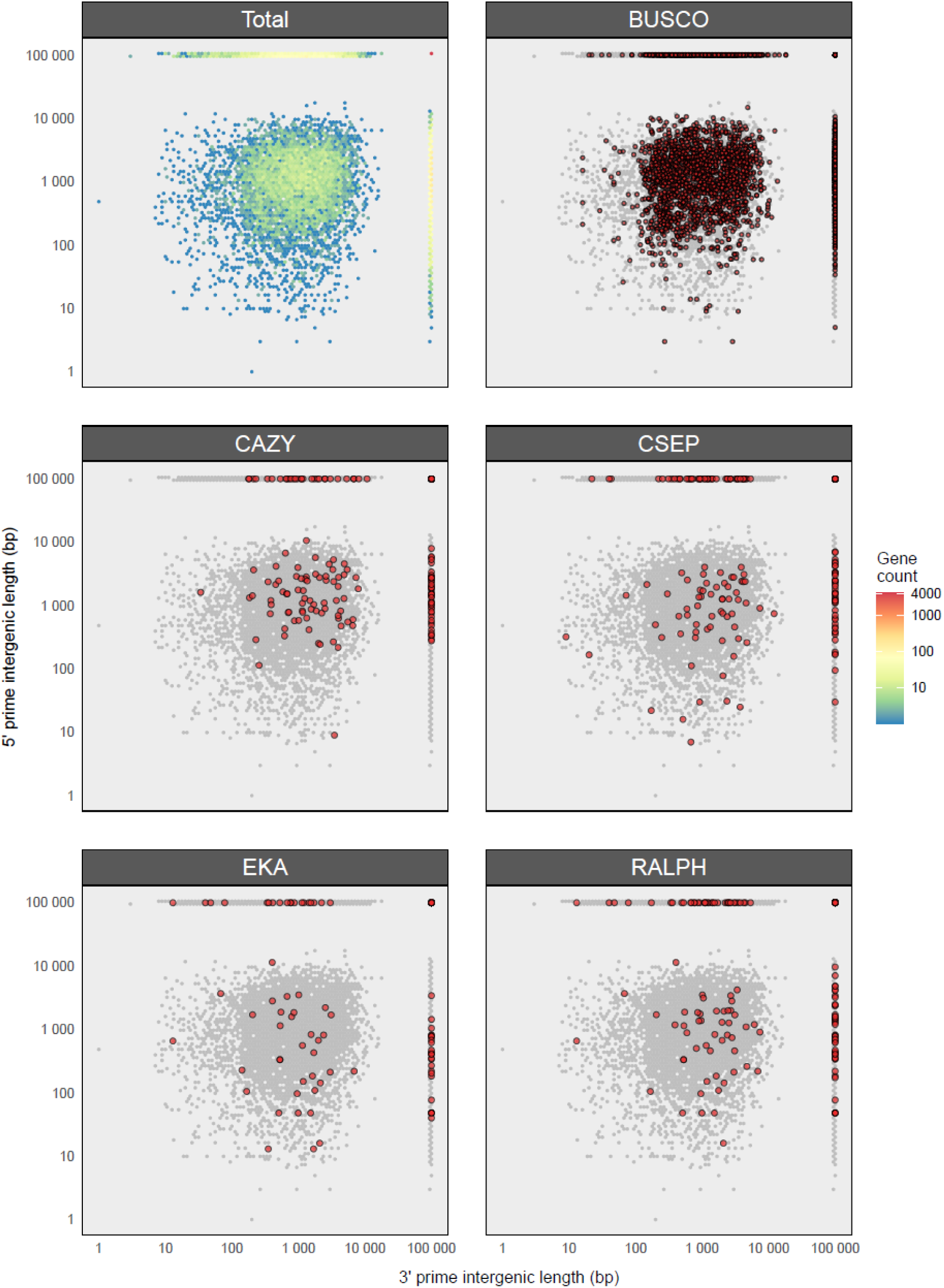
Intergenic distances within *Podosphaera aphanis* assembly DRCT72020: Genes are plotted by five prime (y axis) and three prime (x axis) intergenic distances (bp) with scale bar indicating the gene density of all genes (total) and additional panels highlighting the distribution of Benchmarking Universal Single Copy Orthologue genes (BUSCO), Carbohydrate Active enZYme Genes (CAZY), Candidate Secreted Effector Protein genes (CSEP), genes homologous to effectors homologous to *AVR_k1_* and *AVR_A10_* without canonical secretion signals (EKA), and RnAse-Like Proteins associated with the Haustoria genes (RALPH).

**S5:**
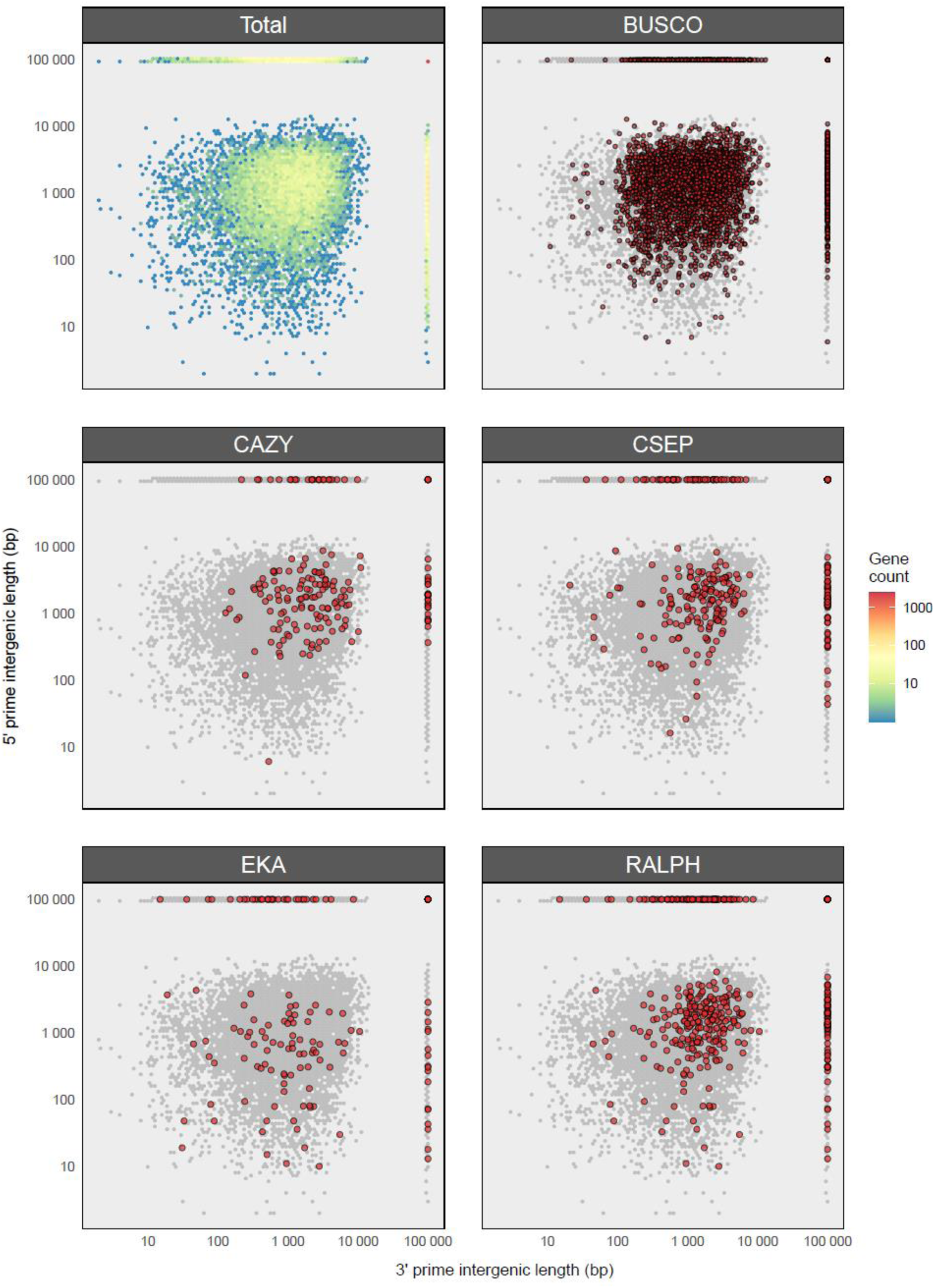
Intergenic distances within *Podosphaera leucotricha* assembly P112020: Genes are plotted by five prime (y axis) and three prime (x axis) intergenic distances (bp) with scale bar indicating the gene density of all genes (total) and additional panels highlighting the distribution of Benchmarking Universal Single Copy Orthologue genes (BUSCO), Carbohydrate Active enZYme Genes (CAZY), Candidate Secreted Effector Protein genes (CSEP), genes homologous to effectors homologous to *AVR_k1_* and *AVR_A10_* without canonical secretion signals (EKA), and RnAse-Like Proteins associated with the Haustoria genes (RALPH).

**S6: Intergenic distance permutation test:** Average three prime, five prime and total intergenic distances are given for Candidate Secreted Effector Proteins (CSEPs), RnAse-Like Proteins associated with the Haustoria (RALPH) homologues, Effectors homologous to AVR_k1_ and AVR_a10_ (EKA), and Carbohydrate Active enZYmes (CAZYs), as well as control Benchmarking Universal Single Copy Orthologue (BUSCO) genes. The observed difference between these counts is reported along with the number of permutations (of 10,000) where a random set of genes had lower intergenic distance than the group of effector candidates.

**S7: *Ab initio* gene predictions and annotations for powdery mildew pathogens:** Contains species, ID, predicted number of genes, Carbohydrate Active enZYmes (CAZY), Pfam annotated proteins (and Pfam virulence annotations), PHI-base effector orthologs, validated effector orthologs, canonically (SP) and Non-Canonically Secreted Proteins (NCSP). Numbers of Candidate Secreted Effector Proteins (CSEP), EffectorP3 effector predictions, and Small Secreted Cysteine-rich Proteins (SSCP) are given, and in brackets the number without orthologues in PHI-base or Predector’s effector database. Finally, the number of candidate effector identifies by either validated effector orthology or *de novo* by secretion signal and EffectorP3 is given.

**S8.**
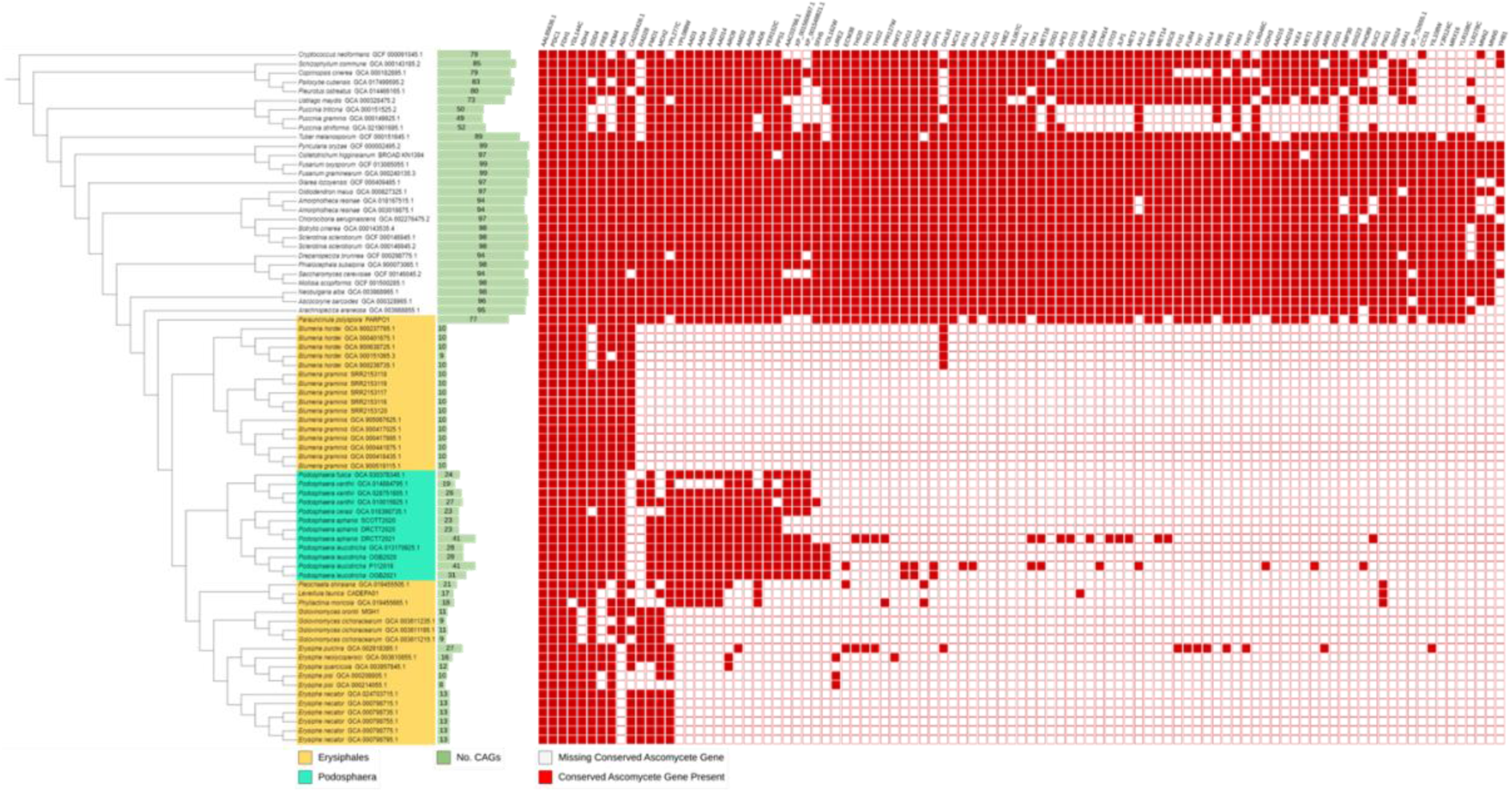
Erysiphaceae genera demonstrate distinct patterns of loss of Missing Conserved Ascomycete Genes (MCAGs): A phylogeny based upon Leotiomycetes benchmarking universal single copy orthologues is presented with members of the mildew family (Erysiphaceae) highlighted in yellow and members of the *Podosphaera* genus highlighted in cyan, alongside this the number of MCAGs with orthologues in the predicted proteome for each assembly is given in green, followed by presence (red) or absence (white) for each MCAG.

**S9: *Podosphaera leucotricha*- and *Podosphaera aphanis*-exclusive and consensus proteome annotations**: presented are gene annotations for the assembled apple powdery mildew genome P112020 and strawberry powdery mildew genome DRCT72020, differentiated by whether given predictions are within orthogroups common to both *P. aphanis* and *P. leucotricha* or exclusive to one species or the other. Included are: the number of gene predictions; the number of resulting proteins with annotations in the Pfam database; with homology to a database of previously-characterised effector genes supplied in the Predector pipeline including both EKA and RALPH homologues. Predicted numbers of Secreted Proteins (SPs) and Candidate Secreted Effector Proteins (CSEP) with EffectorP3 predictions of effectors are given, and in brackets, the number of these proteins which have no hits in PHI-base or the collection of known effector proteins. Finally the number of putative effectors from all annotation methods.

**S10: Apple and strawberry powdery mildew orthologous gene annotations:** gene annotations for the apple powdery mildew assembly P112020 and strawberry powdery mildew assembly DRCT72020 are presented, differentiated by whether given genes are within orthogroups common to both strawberry mildew and apple mildew assemblies or exclusive to one or the other. The raspberry powdery mildew genome, SCOTT2020, was not included in this analysis. Included are: the number of gene predictions; the number of resulting proteins with annotations in the Pfam database; with homology to a database of previously-characterised effector genes supplied in the Predector pipeline including both EKA and RALPH homologues. Predicted numbers of Secreted Proteins (SPs) and Candidate Secreted Effector Proteins (CSEP) with EffectorP3 predictions of effectors are given, and in brackets, the number of these proteins which have no hits in PHI-base or the collection of known effector proteins. Finally the number of putative effectors from all annotation methods.

**S11: *Podosphaera leucotricha*- and *Podosphaera aphanis*-exclusive effector orthologues:** effectors which have been experimentally verified in other species for which orthologues have been identified in the genomes of either *P. leucotricha* or *P. aphanis* are presented. Given is the number of genes in *P. leucotricha* (P112020) and *P. aphanis* (DRCT72020) assemblies that are orthologous to each validated effector and are contained within orthogroups exclusive to that species (Figure 4). Many genes are orthologous to multiple validated effectors. The total number of genes with homology and the number for which a given validated effector had the highest homology (by BLAST E-value) are provided.

